# Transcriptomics-based drug repositioning pipeline identifies therapeutic candidates for COVID-19

**DOI:** 10.1101/2020.10.23.352666

**Authors:** Brian L. Le, Gaia Andreoletti, Tomiko Oskotsky, Albert Vallejo-Gracia, Romel Rosales, Katharine Yu, Idit Kosti, Kristoffer E. Leon, Daniel G. Bunis, Christine Li, G. Renuka Kumar, Kris M. White, Adolfo García-Sastre, Melanie Ott, Marina Sirota

**Affiliations:** Department of Pediatrics, UCSF, SF, CA, USA; Bakar Computational Health Sciences Institute, UCSF, SF, CA, USA; Gladstone Institute of Virology, Gladstone Institutes, SF, CA, USA; Department of Microbiology, Icahn School of Medicine at Mount Sinai, New York, NY, USA; Global Health and Emerging Pathogens Institute, Icahn School of Medicine at Mount Sinai, New York, NY, USA; Biomedical Sciences Graduate Program, UCSF, SF, CA, USA; Shanghai American School, Shanghai, China; Department of Medicine, Division of Infectious Diseases, Icahn School of Medicine at Mount Sinai, New York, NY, USA; The Tisch Cancer Institute, Icahn School of Medicine at Mount Sinai, New York, NY, USA; Department of Medicine, UCSF, SF, CA, USA

**Author notes:** These authors contributed equally to this manuscript (Brian L. Le, Gaia Andreoletti, Tomiko Oskotsky).

## Abstract

The novel SARS-CoV-2 virus emerged in December 2019 and has few effective treatments. We applied a computational drug repositioning pipeline to SARS-CoV-2 differential gene expression signatures derived from publicly available data. We utilized three independent published studies to acquire or generate lists of differentially expressed genes between control and SARS-CoV-2-infected samples. Using a rank-based pattern matching strategy based on the Kolmogorov-Smirnov Statistic, the signatures were queried against drug profiles from Connectivity Map (CMap). We validated sixteen of our top predicted hits in live SARS-CoV-2 antiviral assays in either Calu-3 or 293T-ACE2 cells. Validation experiments in human cell lines showed that 11 of the 16 compounds tested to date (including clofazimine, haloperidol and others) had measurable antiviral activity against SARS-CoV-2. These initial results are encouraging as we continue to work towards a further analysis of these predicted drugs as potential therapeutics for the treatment of COVID-19.

## INTRODUCTION

SARS-CoV-2 has already claimed at least a million lives, has been detected in at least 40 million people, and has likely infected at least another 200 million. The spectrum of disease caused by the virus can be broad ranging from silent infection to lethal disease, with an estimated infection-fatality ratio around 1%^1^. SARS-CoV-2 infection has been shown to affect many organs of the body in addition to the lungs^2^. Three epidemiological factors increase the risk of disease severity: increasing age, decade-by-decade, after the age of 50 years; being male; and various underlying medical conditions^1^. However, even taking these factors into account, there is immense interindividual clinical variability in each demographic category considered^3^. Recently, researchers found that more than 10% of people who develop severe COVID-19 have misguided antibodies―autoantibodies―that attack the innate immune system. Another 3.5% or more of people who develop severe COVID-19 carry specific genetic mutations that impact innate immunity. Consequently, both groups lack effective innate immune responses that depend on type interferon, demonstrating a crucial role for type interferon in protecting cells and the body from COVID-19. Whether the type interferon has been neutralized by autoantibodies or―because of a faulty gene―is produced in insufficient amounts or induced an inadequate antiviral response, the absence of type IFN-mediated immune response appears to be a commonality among a subgroup of people who suffer from life-threatening COVID-19 pneumonia^3^.

While numerous efforts are underway to identify potential therapies targeting various aspects of the disease, there is a paucity of clinically proven treatments for COVID-19. There have been efforts to therapeutically target the hyperinflammation associated with severe COVID-19^4^, as well as to utilize previously identified antiviral medications^5,6^. One of these antivirals, remdesivir, an intravenously administered RNA-dependent RNA polymerase inhibitor, showed positive preliminary results in patients with severe COVID-19^7^. In October 2020, the FDA approved remdesivir for the treatment of COVID-19^8^. Dexamethasone has also been shown to reduce the mortality rate in cases of severe COVID-19^9^.

Nevertheless, the lack of treatments and the severity of the current health pandemic warrant the exploration of rapid identification methods of preventive and therapeutic strategies from every angle. The traditional paradigm of drug discovery is generally regarded as protracted and costly, taking approximately 15 years and over $1 billion to develop and bring a novel drug to market^10^. The repositioning of drugs already approved for human use mitigates the costs and risks associated with early stages of drug development, and offers shorter routes to approval for therapeutic indications. Successful examples of drug repositioning include the indication of thalidomide for severe erythema nodosum leprosum and retinoic acid for acute promyelocytic leukemia^11^. The development and availability of large-scale genomic, transcriptomic, and other molecular profiling technologies and publicly available databases, in combination with the deployment of the network concept of drug targets and the power of phenotypic screening, provide an unprecedented opportunity to advance rational drug design.

Drug repositioning is being extensively explored for COVID-19. High-throughput screening pipelines have been implemented in order to quickly test drug candidates as they are identified^12–15^. In the past, our group has successfully applied a transcriptomics-based computational drug repositioning pipeline to identify novel therapeutic uses for existing drugs^16^. This pipeline leverages transcriptomic data to perform a pattern-matching search between diseases and drugs. The underlying hypothesis is that for a given disease signature consisting of a set of up and down-regulated genes, if there is a drug profile where those same sets of genes are instead down-regulated and up-regulated, respectively, then that drug could be therapeutic for the disease. This method has shown promising results for a variety of different indications, including inflammatory bowel disease^17^, dermatomyositis^18^, cancer^19–21^, and preterm birth^22^.

In existing work from Xing et al.^23^, this pipeline has been used to identify potential drug hits from multiple input disease signatures derived from SARS-CoV or MERS-CoV data. The results were aggregated to obtain a consensus ranking, with 10 drugs selected for *in vitro* testing against SARS-CoV-2 in Vero E6 cell lines, with four drugs (bortezomib, dactolisib, alvocidib and methotrexate) showing viral inhibition^23^. However, this pipeline has not yet been applied specifically to SARS-CoV-2 infection.

A variety of different transcriptomic datasets related to SARS-CoV-2 were published in the spring of 2020. In May 2020, Blanco-Melo et al. studied the transcriptomic signature of SARS-CoV-2 in a variety of different systems, including human cell lines and a ferret model^24^. By infecting human adenocarcinomic alveolar basal epithelial cells with SARS-CoV-2 and comparing to controls, the authors generated a list of 120 differentially expressed genes. They observed two enriched pathways: one composed primarily of type-I interferon-stimulated genes (ISGs) involved in the cellular response to viral infection; and a second composed of chemokines, cytokines, and complement proteins involved in the humoral response. After infecting the cell lines, Blanco-Melo et al. did not detect either ACE2 or TMPRSS2, which are the SARS-CoV-2 receptor and SARS-CoV-2 protease, respectively^25^. However, supported viral replication was observed, thereby allowing the capture of some of the biological responses to SARS-CoV-2.

In May 2020, another study by Lamers et al. examined SARS-CoV-2 infection in human small intestinal organoids grown from primary gut epithelial stem cells. The organoids were exposed to SARS-CoV-2 and grown in various conditions, including Wnt-high expansion media. Enterocytes were readily infected by the virus, and RNA sequencing revealed upregulation of cytokines and genes related to type I and III interferon responses^26^.

A limited amount of transcriptomic data from human samples has also been published. One study detailed the transcriptional signature of bronchoalveolar lavage fluid (of which responding immune cells are often a primary component) of COVID-19 patients compared to controls^27^. Despite a limited number of samples, the results were striking enough to reveal inflammatory cytokine profiles in the COVID-19 cases, along with enrichments in the activation of apoptosis and the P53 signaling pathways.

On the drug side, data are available in the form of differential gene expression profiles from testing on human cells. Publicly-available versions include the Connectivity Map (CMap)^28^, which contains genome-wide testing on approximately 1,300 drugs, wherein the differential profile for a drug was generated by comparing cultured cells treated with the drug to untreated control cultures.

Here, we applied our existing computational drug repositioning pipeline to identify drug profiles with significantly reversed differential gene expression compared to several diverse input signatures for SARS-CoV-2 effects on human cells. By taking into account a broader view of differentially expressed gene sets from both cell line and organoid disease models and human samples, the predictions are complementary to other drug discovery approaches. We identified 102 unique drug hits, from which 25 were identified in at least two of the signatures, several of which have been already investigated in clinical trials. We furthermore explore our findings in the context of other computational drug repurposing efforts for COVID-19. Finally, we tested 16 of our top predicted hits in live SARS-CoV-2 antiviral assays. Four of the top predicted inhibitors were tested for virus inhibition in a human lung cell line, Calu-3, infected with SARS-CoV-2 with quantitation of the secreted virus assessed by RT-qPCR assay. Thirteen predicted inhibitors (including one tested in Calu-3) were incubated with SARS-CoV-2 infected human embryonic kidney 293T cells overexpressing ACE2 (293T-ACE2) with viral replication determined using an immunofluorescence-based assay.

## RESULTS

In this study, we applied our drug repositioning pipeline to SARS-CoV-2 differential gene expression signatures derived from publicly available RNA-seq data (Figure 1). The transcriptomic data were generated from distinct types of tissues, so rather than aggregating them together, we predicted therapeutics for each signature and then combined the results. We utilized three independent gene expression signatures (labelled “ALV”, “EXP”, and “BALF”), each of which consisted of lists of differentially expressed genes between SARS-CoV-2 samples and their respective controls. The ALV signature was generated from human adenocarcinomic alveolar basal epithelial cells by comparing SARS-CoV-2 infection to mock-infection conditions^24^. The EXP signature originated from a study where organoids, grown from human intestinal cells expanded in Wnt-high expansion media, were infected with SARS-CoV-2 and then compared to controls^26^. The BALF signature was from a contrast of primary human BALF samples from two COVID-19 patients versus three controls^27^. Each of these signatures was contrasted with drug profiles of differential gene expression from CMap.

For each of the input signatures, we applied a significance threshold false discovery rate (FDR) < 0.05. We further applied minimum fold change thresholds in order to identify the driving genes. The ALV signature had only 120 genes, with 109 genes shared with the drug profiles; in order to maintain at least 100 genes for the pattern-matching algorithm to work with, we applied no fold-change threshold. For the EXP signature, we applied a |log_2_FC| > 2 cutoff, resulting in 125 genes for the expansion signature (108 shared with the drug profiles). For the BALF signature, we processed the raw read count data to calculate differential gene expression values. We applied a |log_2_FC| > 4 cutoff, with the BALF data yielding 1,349 protein-coding genes for the lavage fluid signature (941 shared with the drug profiles). The gene lists for each of these signatures can be found in the supplement (Tables S1, S2, S3).

We used GSEA (Gene Set Enrichment Analysis)^29,30^ to annotate enriched Hallmark pathways from each of the input signatures (Figure 2A). A number of pathways common to at least two signatures were found. Interferon alpha response and interferon beta response were upregulated in the ALV and EXP signatures. Adipogenesis and cholesterol homeostasis pathways were downregulated in the EXP and BALF signatures. KRAS signaling, and mTORC1 (mammalian target of rapamycin complex 1) signaling were enriched in all three signatures, but not in the same direction, showing the diversity of effects SARS-CoV-2 may have on human cells, and highlighting a need for utilization of diverse profiles as we do in the present study. When we look at the contributing genes within the three signatures (Figure 2B), we found one overlapping upregulated gene - Dickkopf WNT Signaling Pathway Inhibitor 1 (DKK1). We used the publicly available single-cell RNAseq dataset GSE128033^31^ composed of 13 patients (4 healthy, 3 presenting with mild COVID-19 symptoms, and 6 presenting with severe COVID-19 symptoms) to further characterise the expression of DKK1 (Figure S1). Data were re-analyzed following the standard Seurat pipeline. From the analyses of the single-cell data, DKK1 is highly expressed in COVID-19 patients compared to controls, specifically in severe patients and it is expressed by epithelial cells.

After analyzing the input SARS-CoV-2 signatures, we utilized our repositioning pipeline to identify drugs with reversed profiles from CMap (Figure 1). Significantly reversed drug profiles were identified for each of the signatures using a permutation approach: 30 hits from the ALV signature (Table S4), 15 hits from the EXP signature (Table S5), and 86 hits from the BALF signature (Table S6). When visualizing the gene regulation of the input signatures and their respective top 15 drug hits, the overall reversal pattern can be observed (Figure 2C-E). In total, our analysis identified 102 unique drug hits (Table S7). Twenty-five common drug hits were shared by at least two of the signatures (p = 0.0334), with four consensus drug hits (bacampicillin, clofazimine, haloperidol, valproic acid) across all three signatures (p = 0.0599) (Table 1, Figure 3A).

**Table 1.** Therapeutic hits reversing at least 2 of input SARS-CoV-2 signatures. A wide variety of drugs were identified by the analysis of multiple signatures. Drug reversal scores are normalized for each signature; drug entries marked “n.s.” were not significant for reversing that signature.

| **Drug hit** | **Description (current uses)** | **ALV Reversal Score** | **EXP Reversal Score** | **BALF Reversal Score** |
| --- | --- | --- | --- | --- |
| Bacampicillin | Antibiotic | 0.789 | 0.790 | 0.596 |
| Benzocaine | Anesthetic | n.s. | 0.766 | 0.546 |
| Ciclopirox | Antifungal | n.s. | 1 | 0.361 |
| Ciclosporin | Immunosuppressant (RA, psoriasis, Crohn's) | 0.756 | n.s. | 0.409 |
| Clofazimine | Antimycobacterial (leprosy) | 0.946 | 0.893 | 0.558 |
| Co-dergocrine mesilate | Ergoid mesylate (dementia, Alzheimer's, stroke) | 0.775 | n.s. | 0.553 |
| Dicycloverine | Antispasmodic (IBS) | 0.847 | n.s. | 0.461 |
| Fludrocortisone | Corticosteroid | n.s. | 0.782 | 0.519 |
| Fluticasone | Steroid (asthma, COPD) | 0.790 | n.s. | 0.463 |
| Haloperidol | Antipsychotic (schizophrenia) | 0.937 | 0.773 | 0.507 |
| Isoxicam | NSAID | n.s. | 0.873 | 0.410 |
| Lansoprazole | Proton-pump inhibitor (acid reflux) | 0.856 | n.s. | 0.370 |
| Levopropoxyphene | Antitussive | n.s. | 0.835 | 0.770 |
| Lomustine | Antineoplastic (Hodgkin's disease, brain tumors) | 0.748 | n.s. | 0.338 |
| Metixene | Anticholinergic (Parkinson's) | 0.759 | n.s. | 0.344 |
| Myricetin | Flavonoid | n.s. | 0.823 | 0.603 |
| Niclosamide | Anthelmintic (tapeworms) | 0.812 | n.s. | 0.360 |
| Nocodazole | Antineoplastic | 0.766 | n.s. | 0.439 |
| Pentoxifylline | Vasodilatory and anti-inflammatory (claudication) | n.s. | 0.791 | 0.552 |
| Sirolimus | Immunosuppressive | n.s. | 0.768 | 0.729 |
| Thiamazole | Antithyroid agent (Graves disease) | n.s. | 0.796 | 0.724 |
| Tocainide | Antiarrhythmic | 0.798 | n.s. | 0.714 |
| Tretinoin | Vitamin A derivative (acne, acute promyelocytic leukemia) | n.s. | 0.854 | 0.579 |
| Valproic acid | Anticonvulsant (seizures, bipolar disorder) | 0.917 | 0.786 | 0.546 |
| Zuclopenthixol | Antipsychotic (schizophrenia) | 0.754 | n.s. | 0.535 |

We further characterized the common hits by examining their interactions with proteins in humans. We used known drug targets from DrugBank^32^ and predicted additional targets using the similarity ensemble approach (SEA)^33^. We visualized the known interactions from DrugBank in a network (Figure 3B). We also aggregated the list of proteins which were found in DrugBank for at least 2 of the common hits (Table S9). The proteins with the most known interactions with our list of 25 drugs included adrenergic receptors (particularly α2 adrenoreceptors), dopamine receptors, and serotonin receptors.

To confirm the validity of our approach, the inhibitory effects of 16 of our drug hits which significantly reversed multiple SARS-CoV-2 profiles were assessed in live antiviral assays. The inhibitory effects of haloperidol, clofazimine, valproic acid, and fluticasone were evaluated in SARS-CoV-2 infected Calu-3 cells (human lung epithelial cell line), with remdesivir also tested as a positive control. From these five, remdesivir and haloperidol inhibited viral replication (Figure 4A), and the inhibitory effect was also observed by microscopy (Figure 4B).

Additionally, 13 drugs (bacampicillin, ciclopirox, ciclosporin, clofazimine, dicycloverine, fludrocortisone, isoxicam, lansoprazole, metixene, myricetin, pentoxifylline, sirolimus, tretinoin) were assessed in a live SARS-CoV-2 antiviral assay. Remdesivir was again used as a positive control. This testing involved six serial dilutions of each drug to inhibit the replication of SARS-CoV-2 in 293T-ACE2 cells using an immunofluorescence-based antiviral assay^34^. All antiviral assays were paired with cytotoxicity assays using identical drug concentrations in uninfected human 293T-ACE2 cells. Positive control remdesivir and 10 of our predicted drugs (bacampicillin, ciclopirox, ciclosporin, clofazimine, dicycloverine, isoxicam, metixene, pentoxifylline, sirolimus, and tretinoin) showed antiviral efficacy against SARS-CoV-2, reducing viral infection by at least 50%, that was distinguishable from their cytotoxicity profile when tested in this cell line (Figure 5). Several inhibitors showed micromolar to sub-micromolar antiviral efficacy, including clofazimine, ciclosporin, ciclopirox, and metixene. These results not only confirm our predictive methods, but have also identified several clinically-approved drugs with potential for repurposing for the treatment of COVID-19.

## DISCUSSION

Here, we used a transcriptomics-based drug repositioning pipeline to predict therapeutic drug hits for three different input SARS-CoV-2 signatures, each of which came from distinct human cell or tissue origins. We found significant overlap of the therapeutic predictions for these signatures. From 102 total drug hits, 25 drugs reversed at least two signatures (p = 0.0334) and 4 drugs reversed all three signatures (p = 0.0599). The diversity of such signatures yet overlap of highlighted drugs underscores the utility of the current pipeline for identification of drugs which might be therapeutic for the diverse effects of SARS-CoV-2 infection.

Twenty-five of our drug hits reversed at least two of the three input signatures (Table 3). Notably, 14 of the 15 hits from the EXP signature were also hits for the BALF signature, despite being generated from different types of tissue. The EXP signature was generated from intestinal tissue, whereas the BALF signature was generated from constituents of the respiratory tract. Among the common hits reversing at least two of the signatures were two immunosuppressants (ciclosporin and sirolimus), an anti-inflammatory medication (isoxicam), and two steroids (fludrocortisone and fluticasone). Sirolimus (or rapamycin), an immunosuppressant and an mTOR inhibitor, is currently undergoing investigation in several clinical trials in COVID-19 patients (NCT04371640, NCT04341675, NCT04461340). Other hits currently in clinical trials for COVID-19 treatment include ciclosporin (NCT04412785, NCT04392531), niclosamide in combination with diltiazem (NCT04558021), and clofazimine in combination with interferon beta-1b (NCT04465695).

Among our four drug hits that reversed all three signatures, three drugs demonstrated in vitro antiviral efficacy - bacampicillin, clofazimine, and haloperidol. Our group found haloperidol decreased viral growth in SARS-CoV-2 infected Calu-3 cells (Figure 4B) in a dose-dependent manner (Figure 4A). Haloperidol is a psychiatric medication that is indicated for the treatment of psychotic disorders including schizophrenia and acute psychosis. By blocking dopamine (mainly D2) receptors in the brain, haloperidol eliminates dopamine neurotransmission which leads to improvement of psychotic symptoms^35^. Haloperidol can also bind to the sigma-1 and sigma-2 receptors, which are implicated in lipid remodeling and cell stress response^12^. As reported by Gordon et al^12^, the SARS-CoV-2 proteins Nsp6 and ORF9c interact with the sigma-1 receptor and the sigma-2 receptor2, respectively. Moreover, they found that haloperidol decreased viral replication in SARS-CoV-2-infected Vero E6 cells. In another more recent study, Gordon et al found in their analysis of a national electronic medical record database that fewer hospitalized COVID-19 patients who were newly prescribed haloperidol and other Sigma-binding typical antipsychotic medications progressed to requiring mechanical ventilation compared to those who were newly prescribed atypical antipsychotic medications that do not bind to Sigma receptors^14^.

Our testing of clofazimine demonstrated submicromolar antiviral effects of this drug in SARS-Co-V-2 infected 293T-ACE2 and Vero E6 cells (Figures 4 and S3). Clofazimine is an orally administered antimycobacterial drug used in the treatment of leprosy. By preferentially binding to mycobacterial DNA, clofazimine disrupts the cell cycle and eventually kills the bacterium^36^. In addition to being an antimycobacterial agent, clofazimine also possesses anti-inflammatory properties primarily by inhibiting T lymphocyte activation and proliferation^37^. Yuan et al. found that clofazimine inhibits SARS-CoV-2 replication by interfering with spike-mediated viral entry and viral RNA replication. Their work also demonstrated that clofazimine has antiviral efficacy against SARS-CoV-2 in human embryonic stem cell-derived cardiomyocytes and in an ex vivo human lung culture system, as well as antiviral synergy with remdesivir demonstrating the potential of clofazimine as part of a combination treatment regimen for COVID-19^38^.

Our group found bacampicillin to have micromolar antiviral efficacy in SARS-CoV-2 infected 293T-ACE2 cells. Bacampicillin is an orally administered prodrug of ampicillin typically prescribed for treating bacterial infections^39^. As identified by SPOKE^40^, bacampicillin was found to downregulate the GDF15 gene and upregulate the NFKB2 (Nuclear Factor Kappa B Subunit 2) gene in studies by CMap^28^ and LINCS^41^. The GDF15 protein acts as a cytokine and is involved in stress response after cellular injury, and the NFKB2 is a central activator of genes involved with inflammation and immune function^42^. Circulating levels of GDF15 have been found to be significantly higher in COVID-19 patients who die^43^. Zhou et al.’s work revealed NF-kappa B signaling as one of the main pathways of coronavirus infections in humans. While the rapid conversion of bacampicillin to ampicillin in vivo makes this prodrug a less optimal therapeutic candidate for COVID-19, our findings nevertheless provide insights into the immunologic and inflammatory landscape from SARS-CoV-2 infection.

Overall, in testing of our drug hits across two human cell line assays, 11 of 16 exhibited inhibition of SARS-CoV-2 infection. In particular, three of our four consensus drug hits demonstrated antiviral efficacy, with haloperidol showing reproducible inhibition in Calu-3 cells, and bacampicillin and clofazimine inhibiting viral activity in 293T-ACE2 cells without cytotoxicity. Many of our tested drugs can be administered orally, and several are on the WHO Model List of Essential medications, including ciclosporin, clofazimine, and haloperidol^44^. These results suggest that our drug repositioning pipeline can rapidly identify readily available potential therapeutics in antiviral contexts.

There are several limitations of our approach that should be recognized. Data generated from cell lines (both the ALV and EXP signatures) might not accurately represent the biological changes and responses in human infection. Moreover, although the BALF signature was generated from fluid recovered from lavage of infected human tissues, this primary response data was aggregated from a very limited sample size (2 cases and 3 controls). Gathering samples from a larger number of patients should generate a more robust gene expression signature and better inform therapeutic predictions. Furthermore, the drug profiles from CMap were generated from cell line data; drug data generated from more relevant tissue cultures (e.g. lung tissue) may generate more appropriate comparisons.

The drug development response for SARS-CoV-2 / COVID-19 is rapidly developing. One drug, remdesivir, recently received FDA approval for the treatment of COVID-19, and numerous other drugs are being actively explored for possible therapeutic value in COVID-19 cases. Utilizing a diverse set of transcriptomic SARS-CoV-2 signatures, our drug repositioning pipeline identified 25 therapeutic candidates. Validation experiments revealed antiviral activity for 11 of 16 drug hits. Further clinical investigation into these drug hits as well as potential combination therapies is warranted.

## METHODS

### Study design

We have previously developed and used a transcriptomics based bioinformatics approach for drug repositioning in various contexts including inflammatory bowel disease, dermatomyositis, and spontaneous preterm birth. For a list of differentially expressed genes, the computational pipeline compares the ranked differential expression of a disease signature with that of a profile^16,19,28^. A reversal score based on the Kolmogorov-Smirnov statistic is generated for each disease-drug pair, with the idea that if the drug profile significantly reverses the disease signature, then the drug could be potentially therapeutic for the disease.

### SARS-CoV-2 gene expression signatures

Blanco-Melo et al. generated a differential gene expression signature using RNA-seq on human adenocarcinomic alveolar basal epithelial cells infected with SARS-CoV-2 propagated from Vero E6 cells (GSE147507)^24^. Due to the fast-moving nature of the research topic, we opted to use this cell line data in lieu of waiting for substantial patient-level data. This work identified 120 differentially expressed genes (DEGs) – 100 upregulated and 20 downregulated. We used these 120 genes as the ALV signature for our computational pipeline (Table S1).

Lamers et al. performed RNA-seq on their organoid samples, from which differentially expressed genes were calculated. These samples were grown in a medium with a Wnt surrogate supplement and infected with SARS-CoV-2 propagated from Vero E6 cells (GSE149312). They detected 434 significant DEGs (FDR < 0.05). We additionally applied a fold-change cutoff (|log_2_ FC| > 2), resulting in 125 genes used as the EXP signature (Table S2).

Xiong et al. performed RNA-seq analysis of BALF samples from two COVID-19 patients (two samples per patient) and three healthy controls. We processed their raw read counts in order to construct a differential signature (see below for details). FASTQ files were downloaded from the Genome Sequence Archive^45,46^ under accession number CRA002390. Paired-end reads were mapped to the hg19 human reference genome using Salmon (v.1.2.0) and assigned Ensembl genes. After read quality control, we obtained quantifications for 55,640 genes in all samples. In order to identify genes differentially expressed between cases and controls for the BALF samples, we quantified gene expression as raw counts. Raw counts were used as inputs to DESeq2 (v.1.24.0 R package) to call differentially expressed genes (DEGs). After adjusting for the sequencing platform, the default settings of DESeq2 were used. Principal components were generated using the DESeq2 function (Figure S2), and heat maps were generated using the Bioconductor package pheatmap (v.1.0.12) using the rlog-transformed counts (Figure S3). Values shown are rlog-transformed and row-normalized. Volcano plots were generated using the Bioconductor package EnhancedVolcano (v.1.2.0) (Figure S4). Retaining only protein-coding genes and applying both a significance threshold and a fold-change cutoff (FDR < 0.05, |log_2_ FC| > 4), we obtained 1,349 genes to be used as the BALF signature (Table S3).

### Pathway enrichment analysis

Functional enrichment gene-set analysis for GSEA (Gene Set Enrichment Analysis) was performed using fgsea (v.1.12.0 R package) and the input gene lists were ranked by log2 fold change. The 50 Hallmark Gene Sets used in the GSEA analysis were downloaded from MSigDB Signatures database^29,47^. For GO (Gene Ontology) terms, identification of enriched biological themes was performed using the DAVID database^48^.

### Drug gene expression profiles

Drug gene expression profiles were sourced from Connectivity Map (CMap), a publicly-available database of drugs tested on cancer cell lines^28^. CMap contains a set of differential gene expression profiles generated from treating cultured human cells with a variety of different drugs and experimental compounds. These profiles were generated using DNA microarrays to assay mRNA expression. These drug profiles are ranked genome-wide profiles (~22,000 genes) of the effects of the drugs on various cell lines. 6,100 gene expression profiles are presented in CMap. A total of 1,309 compounds were tested in up to 5 different cell lines. The overlap between the gene lists of CMap and the SARS-CoV-2 signature is 109 genes.

### Computational gene expression reversal scoring

To compute reversal scores, we used a non-parametric rank-based method similar to the Kolmogorov-Smirnov test statistic. This analysis was originally suggested by the creators of the CMap database and has since been implemented in a variety of different settings^16–19,22,28^. Similar to past works, we applied a pre-filtering step to the CMap profiles to maintain only drug profiles which were significantly correlated with another profile of the same drug. Drugs were assigned reversal scores based on their ranked differential gene expression profile relative to the SARS-CoV-2 ranked differential gene expression signature. A negative reversal score indicated that the drug had a profile which reversed the SARS-CoV-2 signature; that is, up-regulated genes in the SARS-CoV-2 signature were down-regulated in the drug profile and vice versa.

### Statistical analysis

P-values were adjusted using the false discovery rate (FDR; Benjamini-Hochberg) procedure. P-values for individual drug hits were obtained by comparing reversal scores to a distribution of random scores. Negative reversal scores were considered significant if they met the criterion FDR < 0.05. For drugs tested multiple times (e.g. different cell lines), we used the most reversed profile (lowest negative score). For significance values of the number of drugs reversing multiple signatures, we constructed distributions of the common reversal (reversing two of three signatures) and the consensus reversal (reversing three of three signatures) by randomly sampling the same number of drug profiles for each signature from CMap.

### Single-cell data analysis

Quantification files were downloaded from GEO GSE145926. An individual Seurat object for each sample was generated using Seurat v.3. While the data has been filtered by 10x’s algorithm, we still needed to ensure the remaining cells are clean and devoid of artifacts. We calculated three confounders for the dataset: mitochondrial percentage, ribosomal percentage, and cell cycle state information. For each sample, cells were normalized for genes expressed per cell and per total expression, then multiplied by a scale factor of 10,000 and log-transformed. Low quality cells were excluded from our analyses— this was achieved by filtering out cells with greater than 5,000 and fewer than 300 genes and cells with high percentage of mitochondrial and ribosomal genes (greater than 10% for mitochondrial genes, and 50% for ribosomal genes). SCTransform is a relatively new technique that uses “Pearson Residuals” (PR) to normalize the data. PR’s are independent of sequencing depththanks^49^. We “regress out” the effects of mitochondrial and ribosomal genes, and the cell cycling state of each cell, so they do not dominate the downstream signal used for clustering and differential expression. We then performed a lineage auto-update disabled r dimensional reduction (RunPCA function). Then, each sample was merged together into one Seurat object. Data were then re-normalized and dimensionality reduction and significant principal components were used for downstream graph-based, semi-unsupervised clustering into distinct populations (FindClusters function) and uniform manifold approximation and projection (UMAP) dimensionality reduction was used. For clustering, the resolution parameter was approximated based on the number of cells according to Seurat guidelines; a vector of resolution parameters was passed to the FindClusters function and the optimal resolution of 0.8 that established discernible clusters with distinct marker gene expression was selected. We obtained a total of 21 clusters representing the major immune and epithelial cell populations. To identify marker genes driving each cluster, the clusters were compared pairwise for differential gene expression (FindAllMarkers function) using the Likelihood ratio test assuming an underlying negative binomial distribution (negbinom). For visualization of gene expression data between different samples a number of Seurat functions were used: FeaturePlot, VlnPlot and DotPlot.

### Experimental validation

#### Cell Lines

For studies at the Gladstone Institutes, Calu-3 cells, a human lung epithelial cell line (American Type Culture Collection, ATCC HTB-55), were cultured in advanced MEM supplemented with 2.5% fetal bovine serum (FBS) (Gibco, Life Technologies), 1% L-GlutaMax (ThermoFisher), and 1% penicillin/streptomycin (Corning) at 37°C and 5% CO_2_. SARS-CoV-2 Isolate USA-WA1/2020 was purchased from BEI Resources and propagated and titered in Vero E6 cells.

#### Compounds

Selection of compounds for testing was based on side effect profiles and compound availability. Bacampicillin (B0070000), ciclopirox (SML2011-50MG), ciclosporin (C2163000), clofazimine (1138904-200MG), dicycloverine (D1060000), fludrocortisone (1273003-200MG), fluticasone (1285873-100MG), haloperidol (H1512-5G), isoxicam (I1762-1G), lansoprazole (1356916-150MG), metixene (M1808000), myricetin (M6760-10MG), pentoxifylline (1508901-200MG), sirolimus (S-015-1ML), tretinoin (1674004-5X30MG), and valproic acid (1708707-500MG) were purchased from Sigma-Aldrich. Remdesivir (GS-5734) was purchased from Selleckchem.

Compounds were resuspended in DMSO according to manufacturer’s instructions and serially diluted to the relevant concentrations for treatment of infected cells.

#### Infection Experiments

All work involving live SARS-CoV-2 was performed in the BSL3 facility at the Gladstone Institutes with appropriate approvals. Calu-3 cells were seeded in 96-well plates for 24h, infected with SARS-CoV-2 at a multiplicity of infection (MOI) of 0.05, and treated with compounds. 72 hours post infection, supernatant was collected for RNA extraction and the RNA was analyzed using RT-qPCR to quantify viral genomes present in the supernatant. SARS-CoV-2 specific primers targeting the E gene region: 5’-*ACAGGTACGTTAATAGTTAATAGCGT-3’ (Forward) and 5’-ATATTGCAGCAGTACGCACACA-3’ (Reverse)* were used to quantify cDNA on the 7500 Fast Real-Time PCR system (Applied Biosystems). Cells were fixed with paraformaldehyde and used for immunofluorescence analysis with dsRNA antibody (SCICONS) and DAPI stain. Images were acquired and analyzed using ImageXpress Micro Confocal High-Content Imaging System.

#### In Vitro Microneutralization Assay for SARS-CoV-2 Serology and Drug Screening

For studies carried out at Mount Sinai, SARS-CoV-2 was propagated in Vero E6 cells and 293T-ACE2 cells, as previously described in^12,34^. Two thousand cells were seeded into 96-well plates in DMEM (10% FBS) and incubated for 24 h at 37◻°C,5% CO2. Then, 2 h before infection, the medium was replaced with 100 μl of DMEM (2% FBS) containing the compound of interest at concentrations 50% greater than those indicated, including a DMSO control. The Vero E6 cell line used in this study is a kidney cell line; therefore, we cannot exclude that lung cells yield different results for some inhibitors. Plates were then transferred into the Biosafety Level 3 (BSL3) facility and 100 PFU (MOI = 0.025) was added in 50 μl of DMEM (2% FBS), bringing the final compound concentration to those indicated. Plates were then incubated for 48 h at 37◻°C. After infection, supernatants were removed and cells were fixed with 4% formaldehyde for 24 h before being removed from the BSL3 facility. The cells were then immunostained for the viral NP protein (an in-house mAb 1C7, provided by Dr. Thomas Moran) with a DAPI counterstain. Infected cells (488 nM) and total cells (DAPI) were quantified using the Celigo (Nexcelcom) imaging cytometer. Infectivity is measured by the accumulation of viral NP protein in the nucleus of the Vero E6 cells and 293T-ACE2 cells (fluorescence accumulation). Percentage infection was quantified as ((infected cells/total cells) − background) × 100 and the DMSO control was then set to 100% infection for analysis. The IC50 and IC90 for each experiment were determined using the Prism (GraphPad) software. Cytotoxicity was also performed using the MTT assay (Roche), according to the manufacturer’s instructions. Cytotoxicity was performed in uninfected VeroE6 cells with same compound dilutions and concurrent with viral replication assay. All assays were performed in biologically independent triplicates.

## AUTHOR CONTRIBUTIONS

B.L., T.O. and M.S. designed and coordinated the study. B.L. led the drug repurposing efforts. G.A., K.Y., I.K., and C.L. helped with data analyses. SARS-CoV-2 virus assays were led by A.V.G., G.R.K., K.L., R.R., K.W., A.G.S., and M.O. All the authors contributed to making figures, writing and editing the manuscript.

## ACKNOWLEDGEMENTS

This work was funded in part by the Program for Breakthrough Biomedical Research (PBBR) Grant. This research was also partly funded by the Defense Advanced Research Projects Agency (HR0011-19-2-0020); by CRIP (Center for Research for Influenza Pathogenesis), a NIAID supported Center of Excellence for Influenza Research and Surveillance (CEIRS, contract # HHSN272201400008C); by supplements to NIAID grant U19AI135972 and DoD grant W81XWH-20-1-0270; by the generous support of the JPB Foundation and the Open Philanthropy Project (research grant 2020-215611 (5384); and by anonymous donors to A. García-Sastre. M. Ott acknowledges support through a gift from the Roddenberry Foundation and by NIH 5DP1DA038043. We would like to thank members of the Sirota Lab for useful discussion, Dr. Boris Oskotsky for technical support, Edna Rodas for administrative support, Randy Albrecht for support with the BSL3 facility and procedures at the ISMMS, and Richard Cadagan for technical assistance.

## CONFLICT OF INTEREST STATEMENT

M.S. is on the advisory board of twoXAR. The García-Sastre Laboratory has received research support from Pfizer, Senhwa Biosciences and 7Hills Pharma. A.G.S. has consulting agreements for the following companies involving cash and/or stock: Vivaldi Biosciences, Contrafect, 7Hills Pharma, Avimex, Vaxalto, Accurius and Esperovax. Other authors declare no competing financial interests.

---

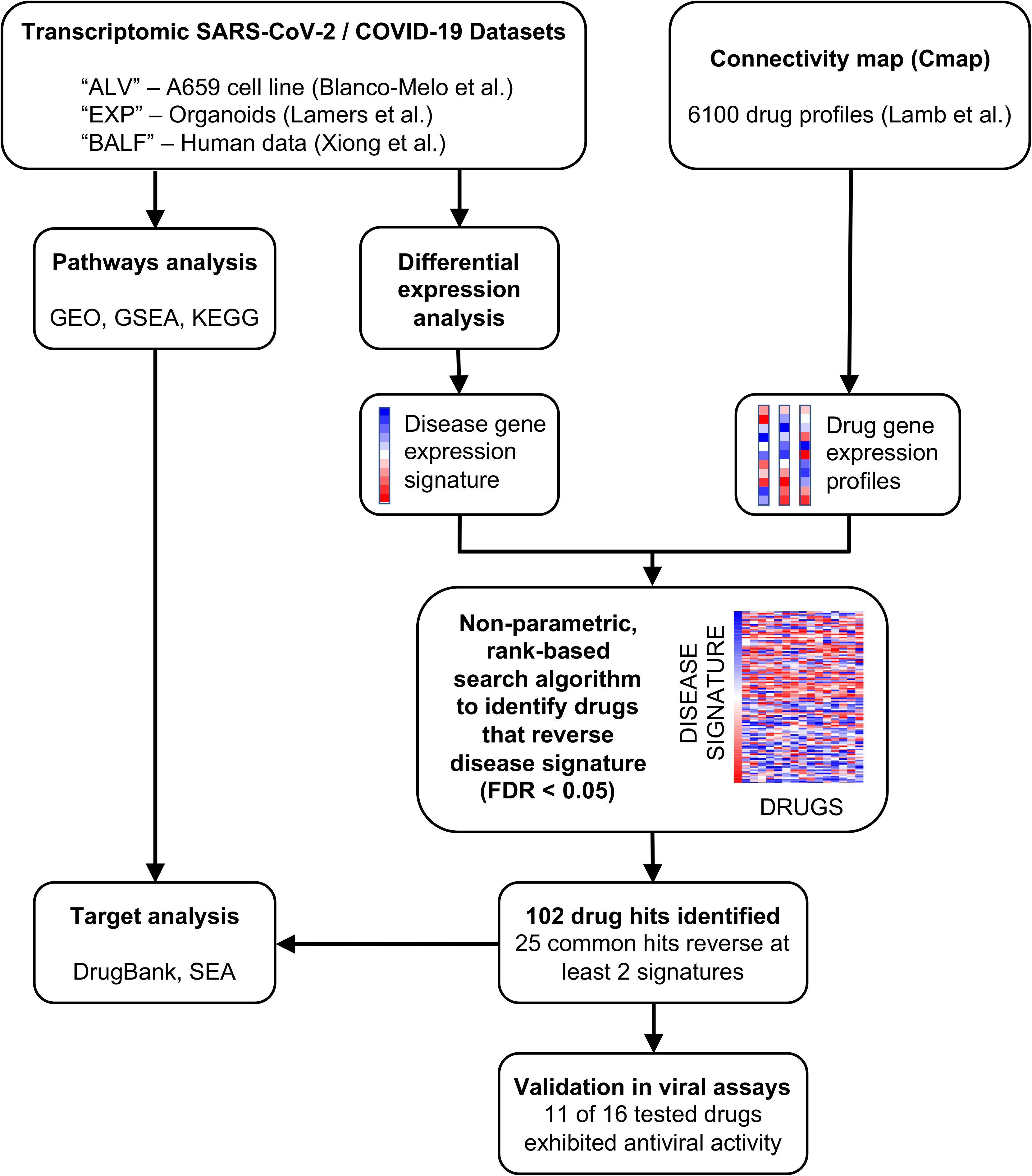

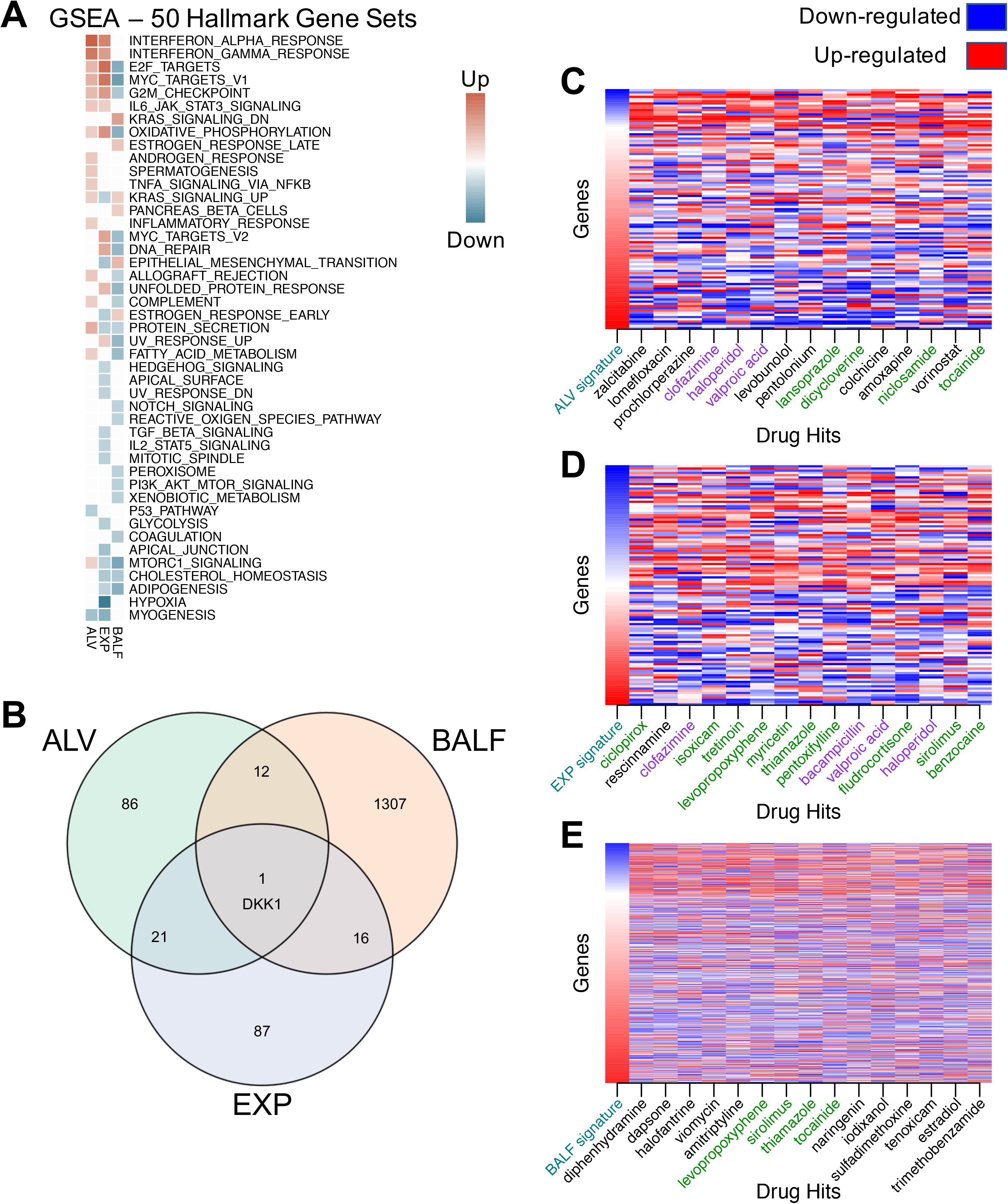

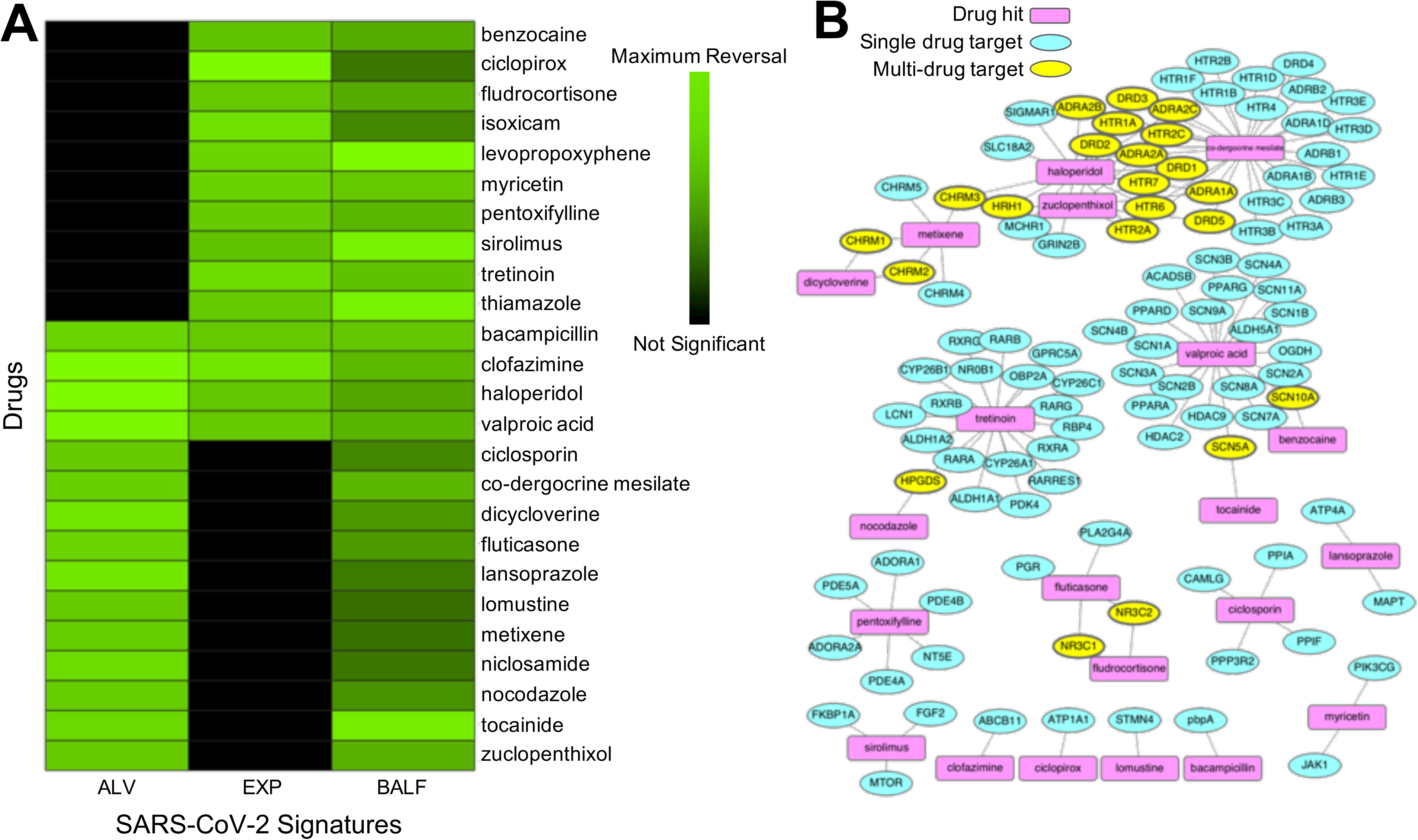

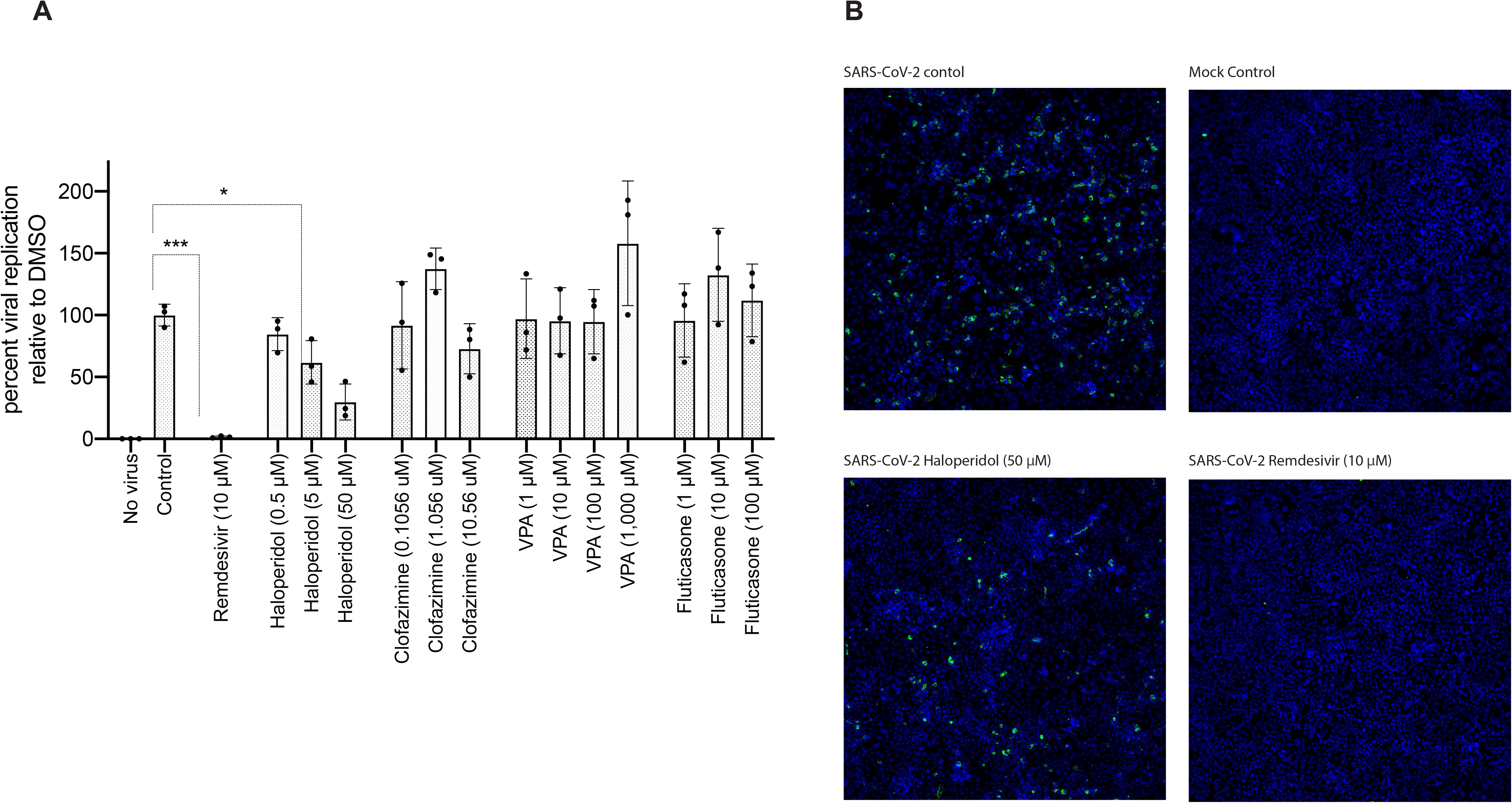

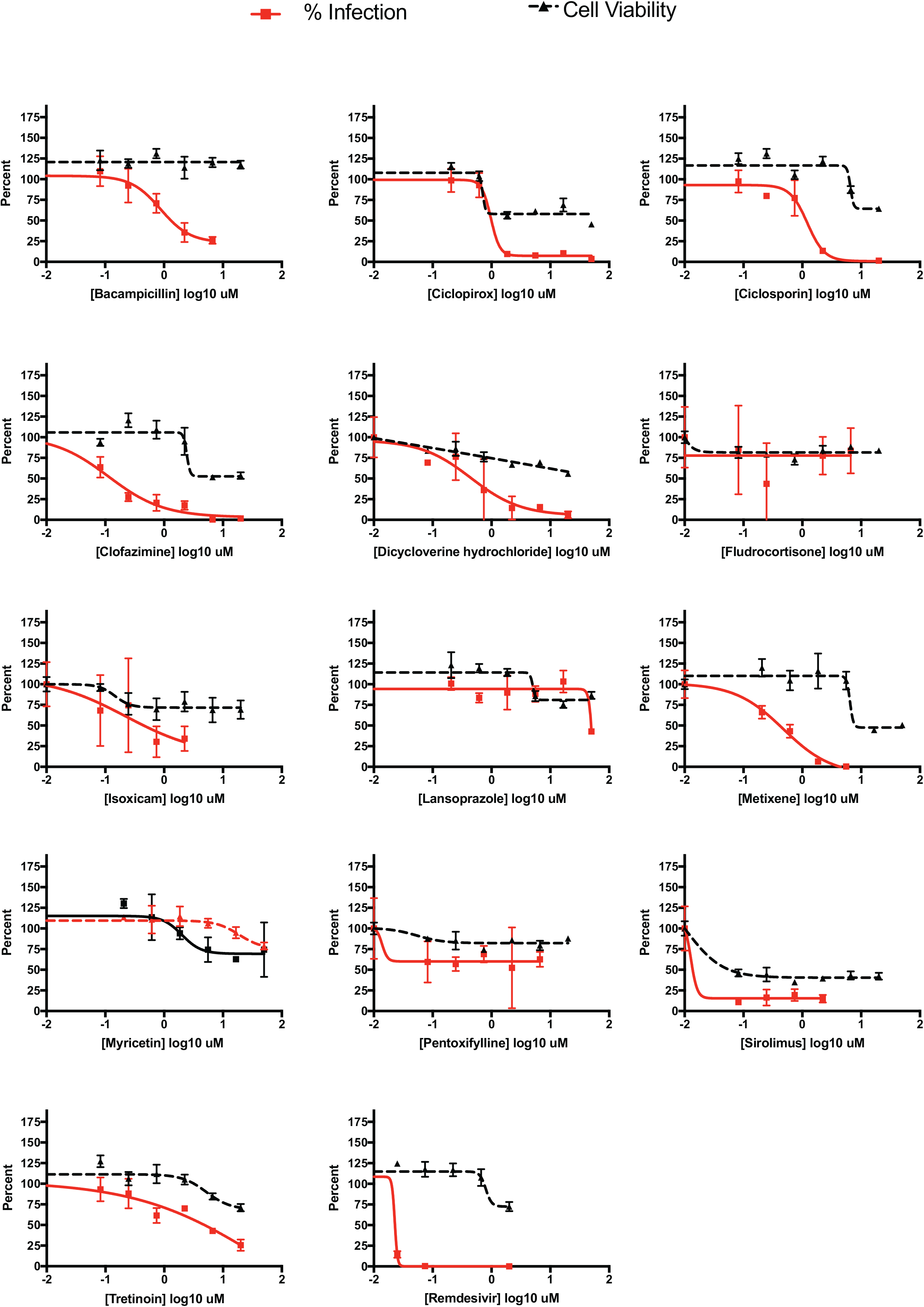

