## Supplementary figures and images for "Transcriptomics-based drug repositioning pipeline identifies therapeutic candidates for COVID-19"

### Supplemental Figure 1

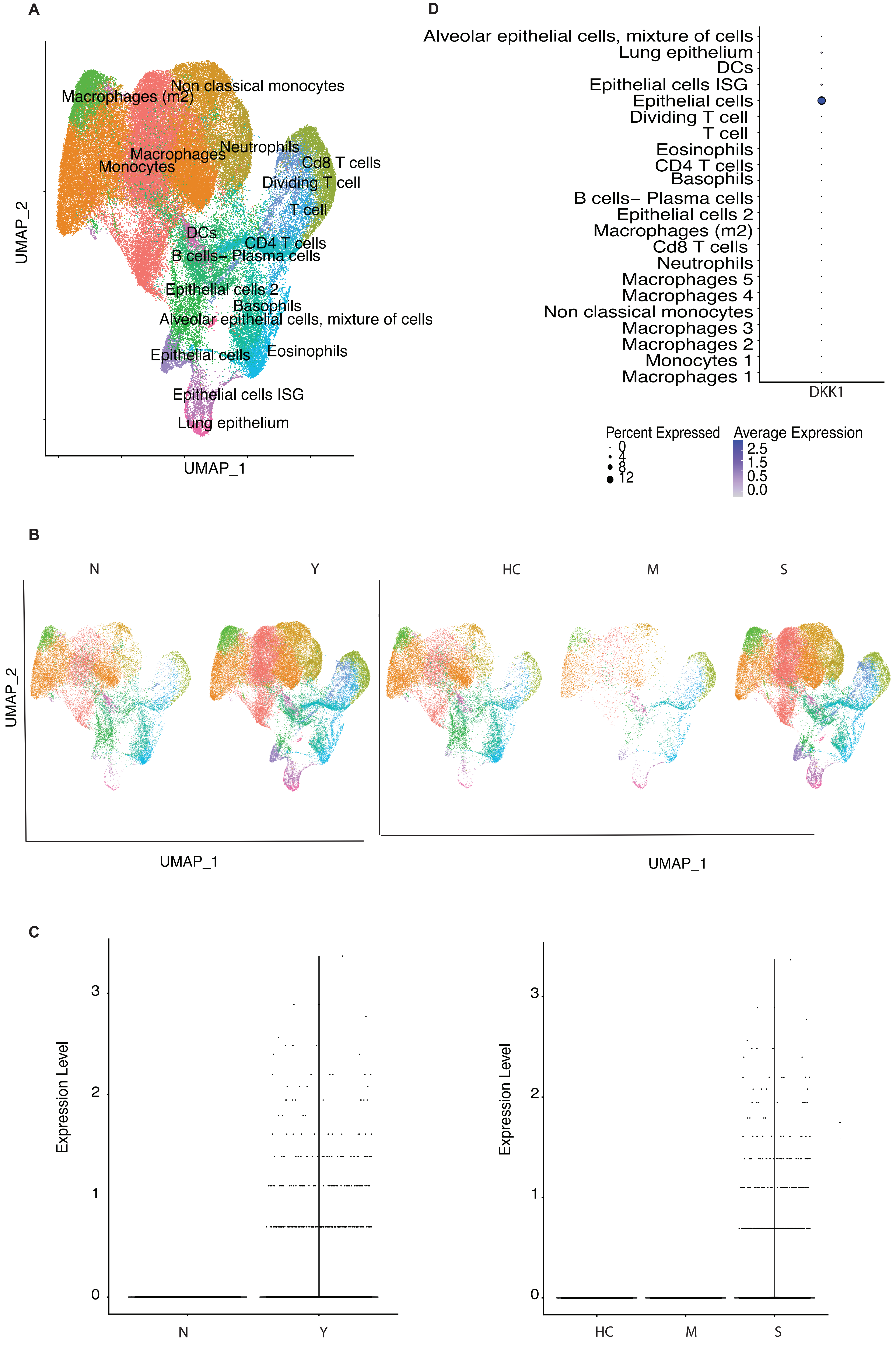

### Supplemental Figure 2

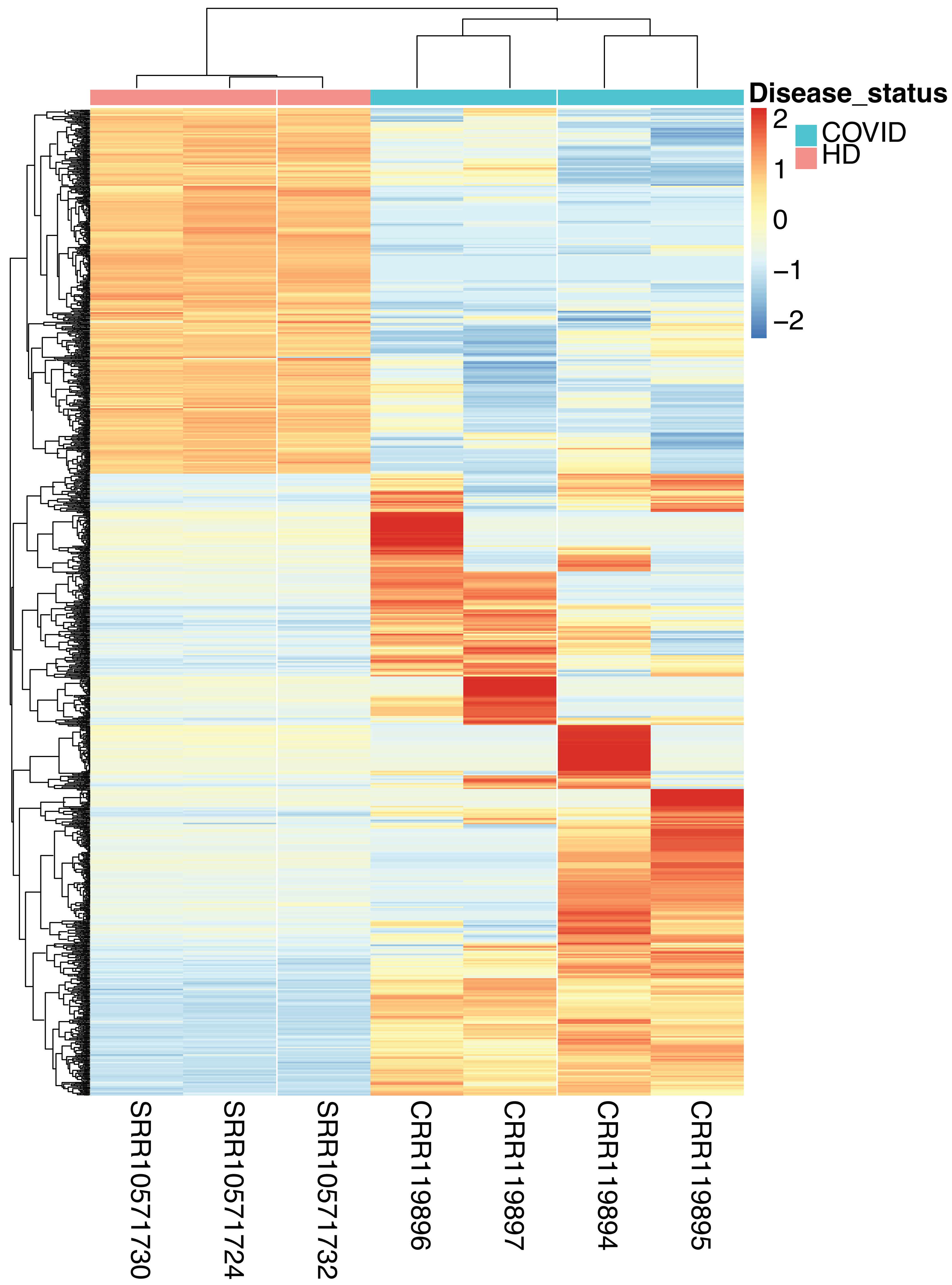

### Supplemental Figure 3

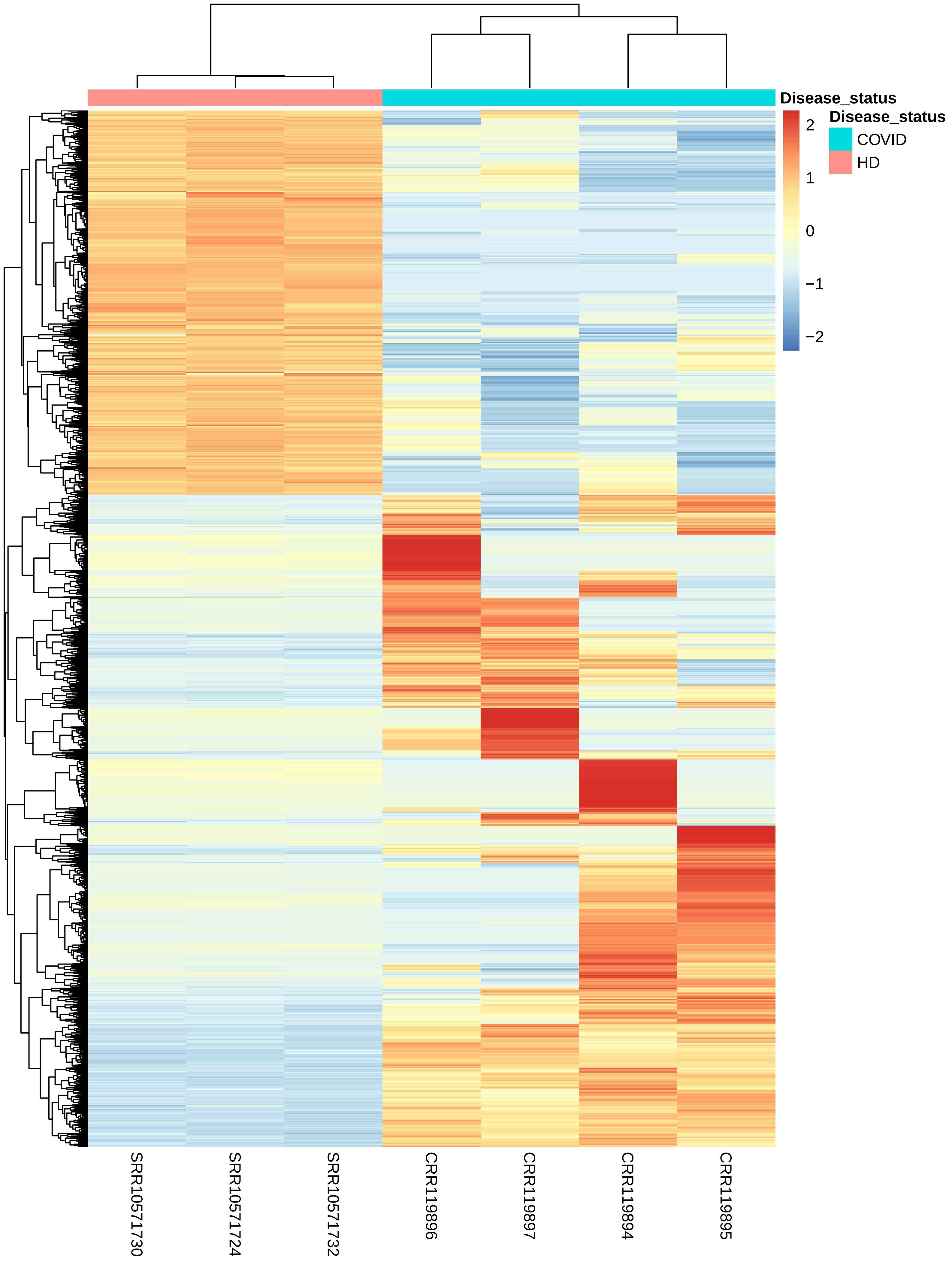

### Supplemental Figure 4

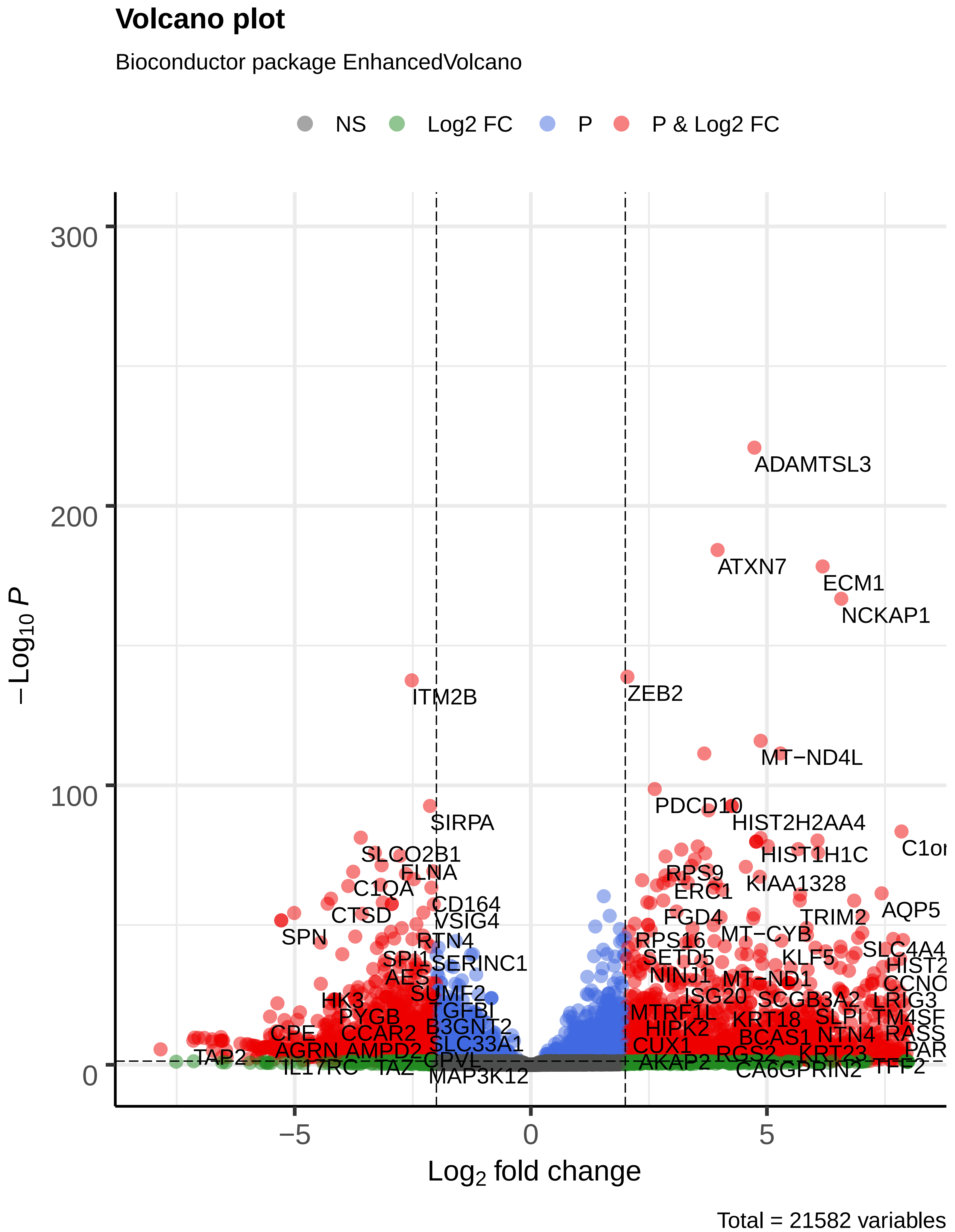

### Supplemental Figure 5

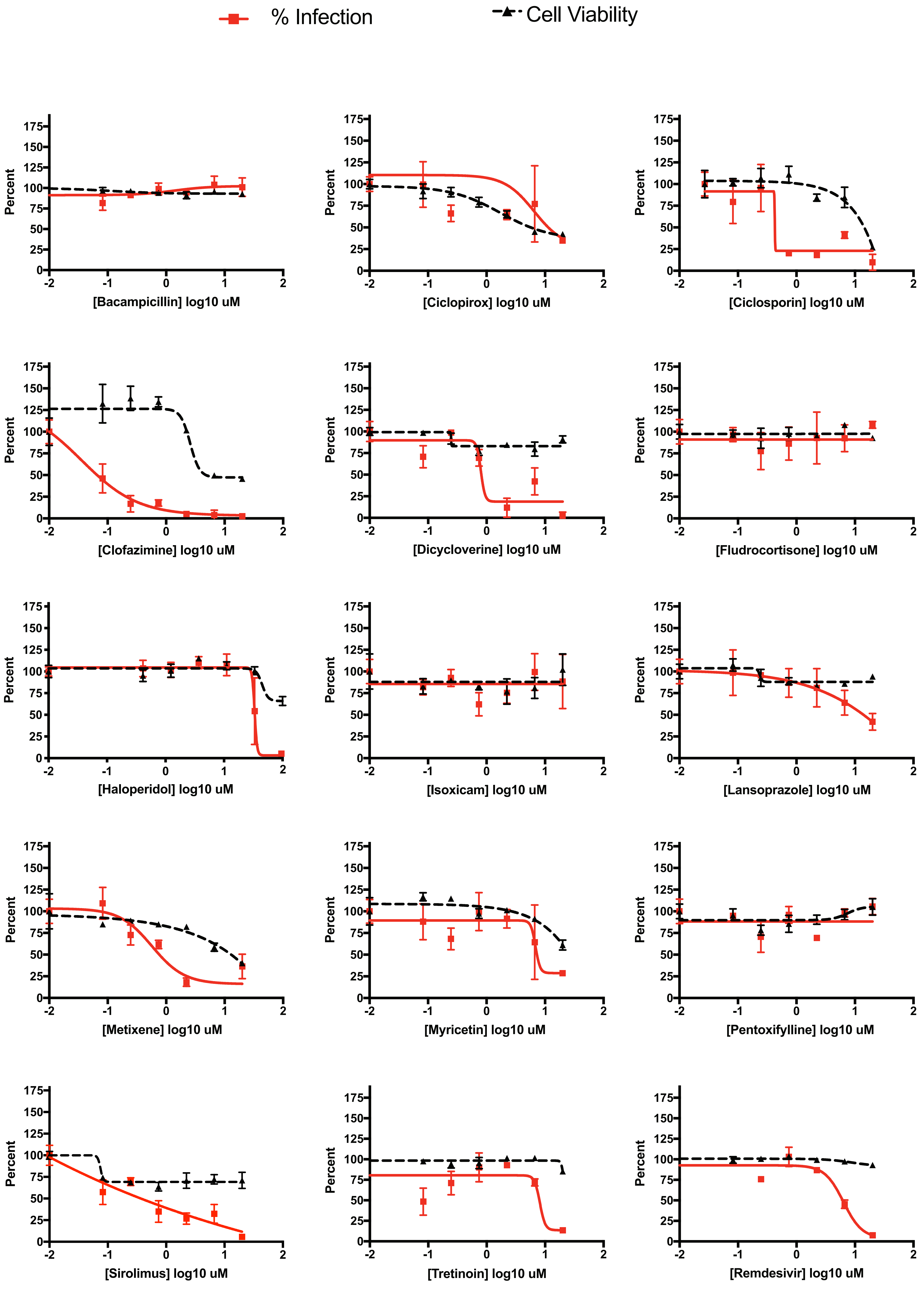
